# Infra-delta oscillatory structure in expressive piano performance: evidence for a shared motor timing mechanism

**DOI:** 10.64898/2026.03.27.714869

**Authors:** Alice Mado Proverbio, Chang Qin

**Affiliations:** Department of Psychology, University of Milano-Bicocca, Milan, Italy; International School of Advanced Studies, University of Camerino (MC), Italy

**Keywords:** Music, Motor timing, Oscillatory dynamics, Music performance, Temporal structure, Sensorimotor synchronization

## Abstract

This study examines the temporal dynamics of expressive piano performance by means of a quantitative analysis of motor timing in an elite pianist, with particular reference to stylistic contrasts between Baroque and Romantic repertoire. In line with kinematic models of expressive timing, which describe musical performance as reflecting principles of biological motion, we examined whether a common temporal structure underlies stylistically divergent executions. Despite marked differences in structural complexity and gesture density, both performances exhibited a shared low-frequency oscillatory pattern (∼0.36 Hz) in beat-level timing variability. This infra-delta rhythmic modulation is consistent with the presence of an underlying motor timing scaffold and suggests a common temporal organization across expressive behaviors. These findings support the hypothesis that musical performance relies on a rhythmically structured control architecture, potentially shared with other complex motor activities such as speech and locomotion.

## Introduction

Tempo and time signature detection constitutes a crucial component of computational music analysis (Kostrzewa & Zabiałowicz, 2023), enabling algorithms to extract rhythmic structure and meter from complex musical signals (Cozens & Godsill, 2024). This technology, particularly beat-tracking, is based on the assumption that no matter how complex, varied and expressive the music is, there exists an underlying, relatively stable temporal framework. Interestingly, contemporary neurobiological models of motor timing are also predicated on this foundational assumption (Mauk, 2004; Ivry & Spencer, 2004; Schwartze and Kotz, 2024). Spontaneous periodic rhythms are evident across a range of human behavioral systems, such as speech (Jungers et al., 2002), locomotion (Murray et al., 1964), and finger tapping (McAuley et al., 2006). Even highly complex behaviors, such as piano performance (Zamm et al., 2018), exhibit spontaneous and relatively stable rhythmic cycles at intrinsic frequencies. Compared to novice pianists, expert performers can sustain precise temporal sequencing and control during performance tasks, even when auditory or sensory feedback is diminished or disrupted (Lappe et al., 2018), by relying on robust internal models and predictive neural mechanisms (Herholz et al., 2012).

Consistently, biomechanical studies have demonstrated that temporal variations in expressive performance often follow parabolic curves (Friberg and Sundberg, 1999; Todd, 1992). For example, the final slowing of a musical piece tends to emulate the natural deceleration of a physical body in motion (motor model). This makes the music sound “natural” to the human ear, as it reflects our embodied experience of gravity and inertia. Models of expressive timing in musical performance can be elegantly described through analogies with physical motion, such as acceleration and deceleration (Clarke, 1988). It appears that the final *ritardando* in musical performance follows a deceleration curve closely resembling that observed in a runner coming to a stop, suggesting that the temporal expressivity of music reflects the dynamics of human movement (Friberg and Sundberg, 1999). Indeed, several studies have demonstrated that piano performance constitutes a highly complex cognitive-motor task, dependent on the brain*’*s internal timing mechanisms to accurately perceive and regulate temporal structures. The central mechanisms underlying this motor control involve the cerebellum and basal ganglia–thalamo–cortical circuitry, which generate predictive temporal signals to facilitate rhythm processing (Parsons et al., 2005; Kasdan et al., 2022; Evers & Stefan, 2023; Kameda et al., 2023; Patel et al., 2014; Vuust & Witek, 2014). The population clock model proposes that temporal perception originates from coordinated group activity in the cerebellum, basal ganglia, and motor cortex (Buonomano & Laje, 2010; Zhou & Buonomano, 2022). The temporal stability observed in piano performance is attributed to dynamic patterns of neural groups rather than individual neuron activity. The oscillator model further posits that neural oscillations shape the perceived temporal structure of complex rhythms (Large et al., 2015), supporting the view that time perception is encoded in dynamic activity of neural populations.

Here, we present a novel system for extracting both motor and musical timing features with high precision from an elite pianist*’*s performance, investigating whether an underlying oscillatory motor structure governs expressive timing. We examined whether motor performance timing, irrespective of musical piece, style, or structure, follows a common hidden temporal framework, potentially reflecting the rhythmic nature of neural signaling.

We deliberately selected two works that differ significantly in historical period, musical style, average tempo, and expressive interpretation: Bach’s Contrapunctus I (BWV 1080) and Chopin’s Ballade No. 1 in G minor (Excerpt). Bach is a representative of the late Baroque period, and his works generally follow strict counterpoint and rhythmic organization (Fabian, 2017); Chopin, one of the founders of romantic piano music, makes extensive use of rubato (free rhythm), complex harmonic progressions and dramatic dynamic contrasts in his works to highlight emotional expression and personalized interpretation (Repp, 1998; 1999). These two works represent the core styles of the Baroque period (approximately 1600–1750) and the Romantic period (approximately 1810–1900), respectively. The contrast between them highlights the universality of the principles of motor control under investigation.

It was hypothesized that, notwithstanding stylistic differences and individual interpretative choices (see Table 1 for a comparison with selected piano recordings), piano performances would nonetheless exhibit a common underlying periodic motor structure.

**Table 1.** Performance details and execution durations for excerpts from J.S. Bach’s Contrapunctus I (BWV 1080) and Frédéric Chopin’s Ballade No. 1 in G minor. The table lists selected interpretations by renowned pianists, indicating the context of each recording (live or studio), venue or label, year of execution, and the total duration of the excerpt analyzed. “Pianist1” refers to the performer recorded during the experimental session. Notably, there is considerable variability in the duration of execution for the same musical excerpts, despite their identical notated content. For Contrapunctus I, performances range from 2’29’’ (Kocsis, 1985) to 4’49’’ (Gould, 2009), revealing almost a twofold difference. Similarly, the Ballade No. 1 excerpt spans from 2’24’’ (Richter, 2012) to 2’44’’ (Barenboim, 2020). These differences likely reflect a combination of interpretative choices, stylistic traditions, and performance contexts—such as tempo flexibility, articulation, phrasing, and rubato—underscoring the expressive latitude permitted by classical performance practice. Recordings were sourced from publicly available content on YouTube.

| Pianist | Execution details | Timing |
| --- | --- | --- |
| Bach's Contrapunctus I (BWV 1080) |  |  |
| András Schiff | Live recording, Shanghai Symphony Hall, 2025 | 2'33 |
| Zoltán Kocsis | Universal International Music B.V., 1985 | 2'29" |
| Daniil Trifonov | Deutsche Grammophon, live recital 2021 | 2'52" |
| Agata Sorotokin | Arthur Fraser International Piano Competition, 2014 | 3'26" |
| **Pianist1** | **Live recording** | **3'35"** |
| Tatiana Nikolayeva | Live recording, Rosslyn Hill Chapel, 1992 | 4'48" |
| Vyacheslav Gryaznov | Live recording, Moscow Conservatory, 2012 | 4'48" |
| Glenn Gould | Firma Melodiya, 2009 | 4'49" |
| Chopin's Ballade No. 1 in G minor (Excerpt) |  |  |
| Sviatoslav Richter | Praga Digitals, 2012 | 2'24" |
| Krystian Zimerman | Deutsche Grammophon GmbH, 1988 | 2'24" |
| Yundi Li | Live recording, Carnegie Hall, 2016 | 2'28" |
| Vladimir Horowitz | Live recording, Royal Festival Hall, London, 1982 | 2'33" |
| Arthur Rubinstein | BNF Collection, 2014. Originally released 1960 | 2'36" |
| Claudio Arrau | Universal International Music B.V., 2001 | 2'37" |
| **Pianist1** | **Live recording** | **2'43"** |
| Daniel Barenboim | Live recording at Pierre Boulez Saal, Berlin, 2020 | 2'44" |

From an applied perspective, the present work contributes to the quantitative characterization of temporal structure in complex motor behaviors, integrating signal analysis, behavioral metrics, and theoretical models of neural timing. This approach aligns with recent efforts in computational neuroscience and performance science to identify low-dimensional dynamical signatures underlying high-level motor expertise.

## Methods

Given the exploratory nature of the study and the focus on high-resolution temporal dynamics, a single-case design was adopted. This approach allows for fine-grained analysis of performance timing while minimizing inter-subject variability, and has been previously employed in studies of expert motor behavior. The use of expert ratings and manual temporal annotation was selected to ensure ecological validity and alignment with performance practice, complementing computational analyses of timing variability.

### Participant

The performer (Pianist1) was a 30 year old, healthy, right-handed elite male pianist with seven years of international concert experience (including Carnegie Hall, Nagoya Philharmonic, and Hong Kong City Hall), fully capable of performing the experimental piano pieces and demonstrating a high level of expressive and interpretative sophistication. Pianist1 began studying music at the age of four, marking the onset of an early and continuous engagement with the piano. The artist has never suffered from any neurological or psychiatric disorders (including motor or cognitive impairments) and has perfectly normal vision and hearing. The participant provided written informed consent, and the research was conducted in accordance with the Declaration of Helsinki and ethical Committee guidelines.

### Procedure and analyses

The pianist performed on a Yamaha P-225B digital piano in an anechoic chamber, utilizing the default piano sound. Audio recordings (WAV format) and behavioral data were collected during the performance. The pianist performed seven pieces over a 30-minute session in a continuous and natural manner without external interference, including Johann Sebastian Bach*’*s The Art of Fugue: Contrapunctus I (BWV 1080), and an excerpt from Frédéric Chopin*’*s Ballade No.1 in G minor, Op. 23). Bach piece was performed based on The Art of Fugue, BWV 1080: Performance Score, edited and engraved by Dr. Dominic Florence, published by Contrapunctus Press in 2021. The Chopin piece was performed using the National Edition of the Works of Fryderyk Chopin, edited by Jan Ekier and Paweł Kamiński, published by PWM Edition. The pianist executed the full program from memory, following recent recordings for a CD and a series of performances on tour.

This study quantified the number of notes and total gestures separately for the left and right hands. Gestures were defined as each new key-press onset event within any passage. The number of notes for each hand was determined by counting the key-press onset events produced independently by the left and right hands. Notes sustained in legato passages, i.e., without new key presses, were excluded from all counts. By combining this data with temporal information extracted from Audacity software, the number of notes and gestures per second for each hand was calculated. Movement complexity was evaluated by two external professional pianists using a six-point Likert scale for each measure. The evaluation criteria included Melodic/Harmonic Complexity, Executive Complexity, and Rhythmic Complexity (Fig. 1). The two pianists subjectively assessed the complexity of the musical structure and the performance difficulty of the passages using the following six-point Likert scale (1–6):

**Fig. 1.**
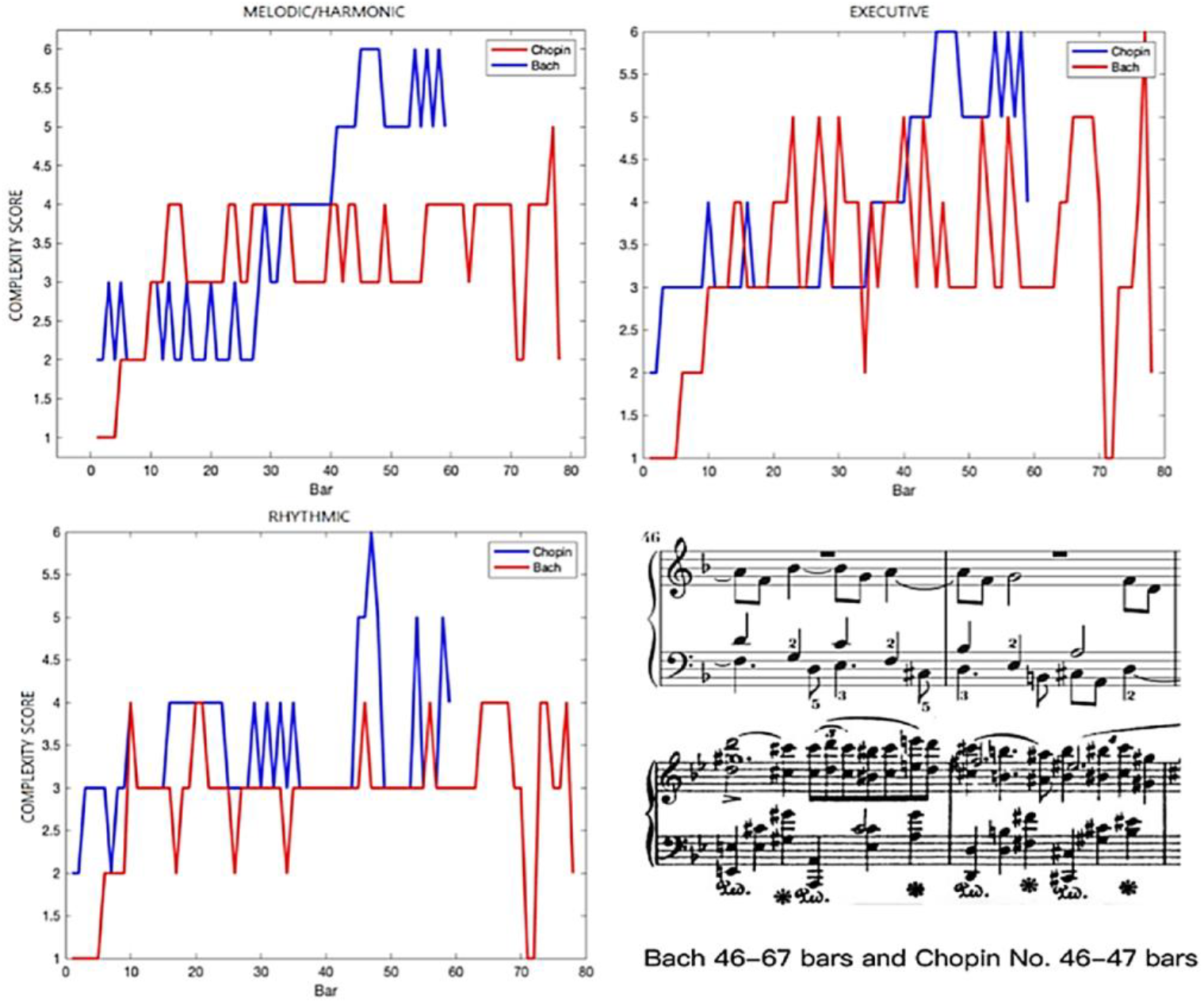
Mean ratings of melodic/harmonic, executive, and rhythmic complexity of the two piano pieces, as assessed by two professional pianists and teachers. In the lower right portion of the figure, two excerpts from the respective pieces are displayed to highlight differences in structural complexity.

1. A single-line melody or an extremely simple harmonic structure; the performance task is very easy to execute.
2. A two-voice melody or a simple three-voice passage, with basic harmonic support; the overall execution is relatively straightforward.
3. A three-voice melody or a simplified four-voice texture; the performance difficulty is moderate.
4. Complex three-voice melodies with interwoven and overlapping voices, or standard four-voice textures, are somewhat challenging to perform.
5. Complex four-voice melodies, or harmonic passages requiring coordination of both hands, are more difficult to perform.
6. Highly complex and variable harmonic textures; the performance tasks are extremely challenging, demanding exceptionally high technical and perceptual skills.

The musical score utilized during the verification process was identical to the one employed by the pianist during the performance. Incomplete measures at the beginning and end of the audio recordings were excluded from the analysis to ensure data consistency. The audio processing was conducted using Audacity software (version 3.7.0). The study employed single-track mode for audio processing, achieving time measurement accuracy at the millisecond level. Beat durations were quantified using the quarter note as the temporal unit of measurement. Beat onset times were determined by identifying the initial rise in the amplitude of the waveform corresponding to each beat, as observed in Audacity. In instances where the left- and right-hand parts did not align temporally as per the musical score, priority was given to the main melodic line for beat identification. Based on the precise timing of each quarter note BPM values (beats per minute, calculated as BPM = 60 / seconds per beat) were calculated per bar (Fig. 2) and per quarter note (Fig. 3). Tactus instability was then assessed both for Contrapunctus I (2/2 tempo) and in Chopin’s Ballade (6/4 tempo).

**Fig. 2.**
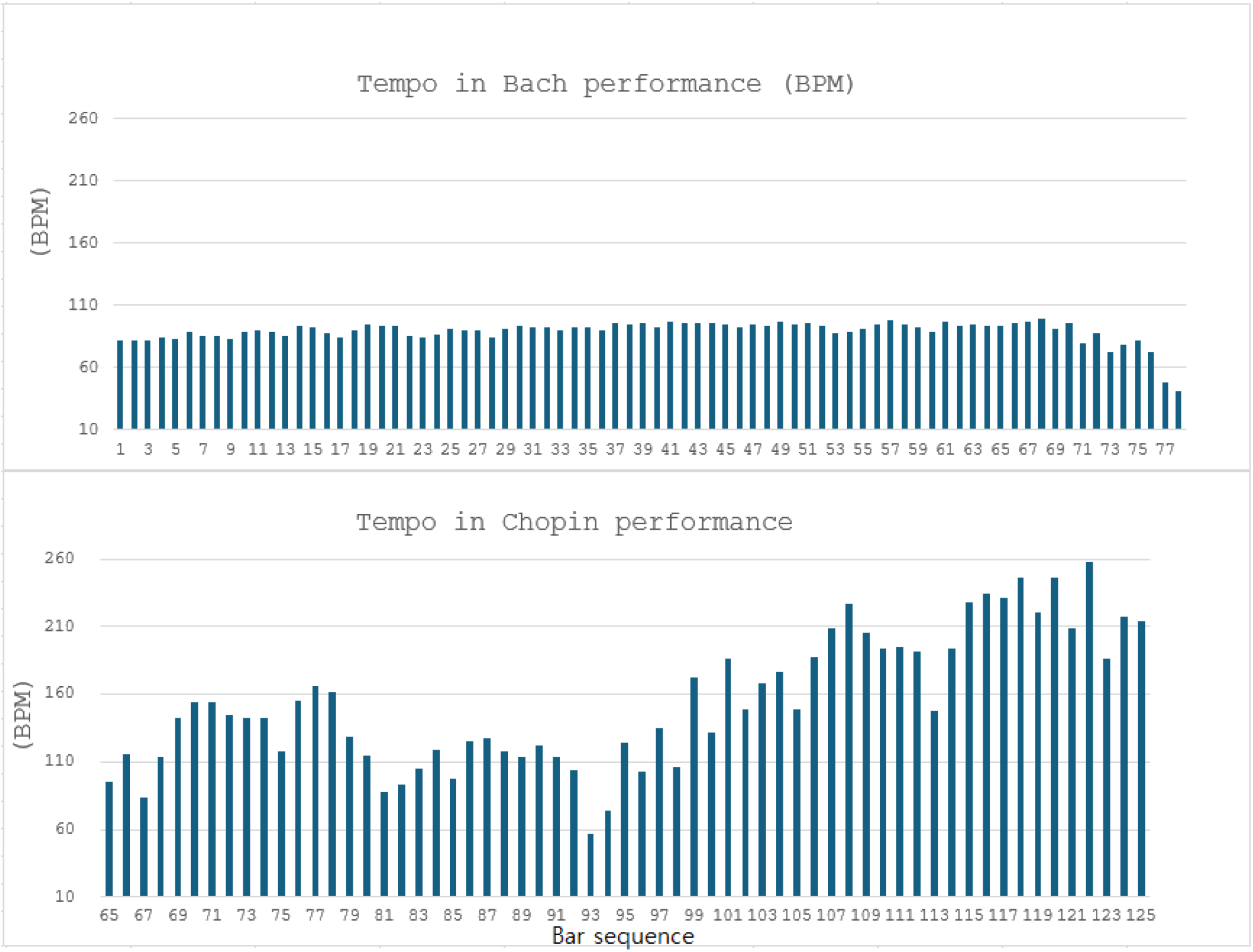
Effective execution tempo (in beats per minute) across individual bars (plotted along the x-axis) in J.S. Bach’s Contrapunctus I, and Chopin’s Ballade no. 1.

**Fig. 3.**
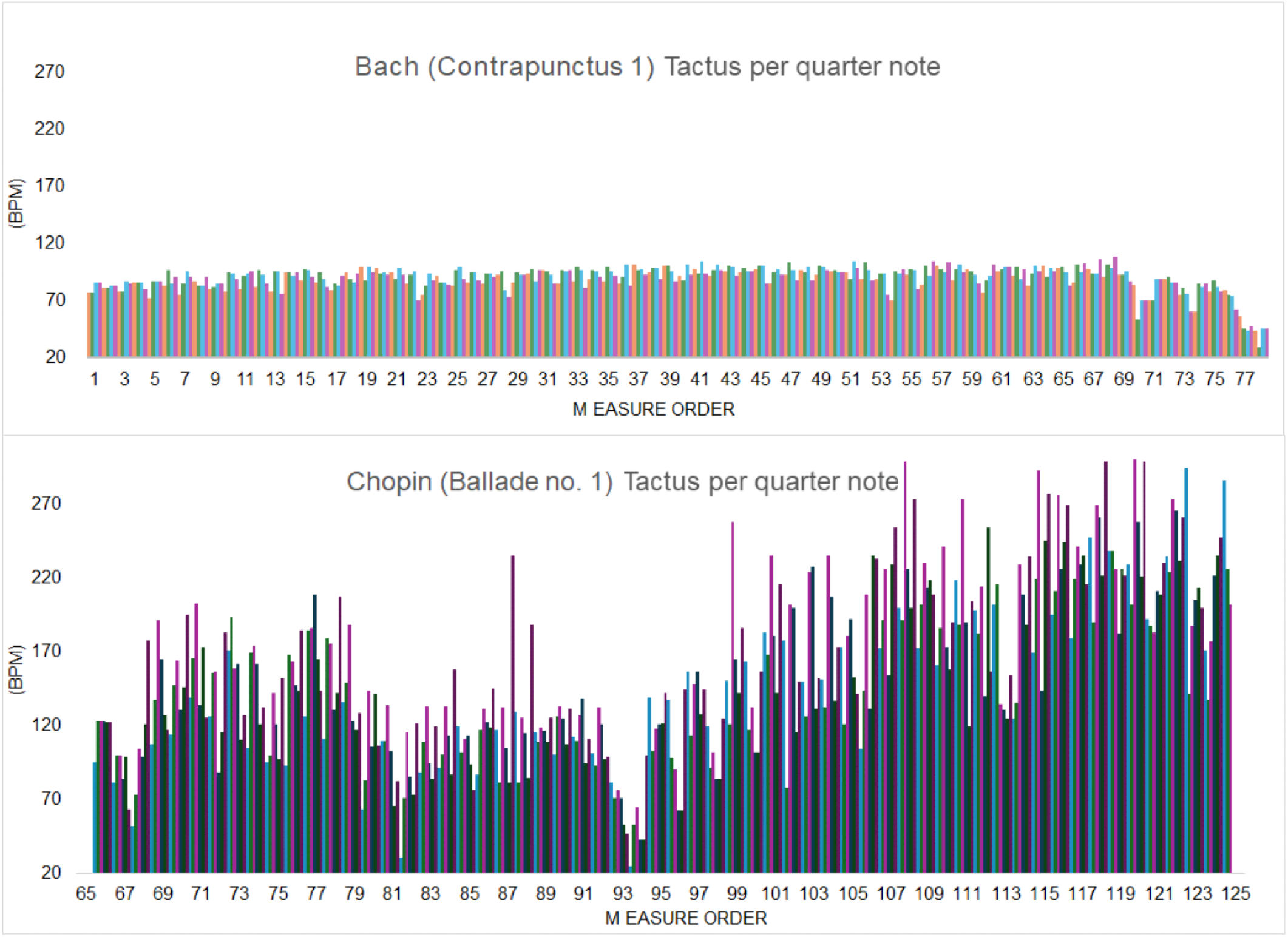
Effective execution tempo (in beats per minute) for each quarter note (plotted along the x-axis) in J.S. Bach’s Contrapunctus I and Chopin’s Ballade No. 1.

### Statistical analyses

To compare the structural complexity of musical passages composed by Chopin and Bach, we conducted non-parametric statistical analyses on a set of quantitative features. Specifically, we extracted the following complexity metrics from each matched excerpt: number of gestures per measure, number of gestures per second, melodic/harmonic complexity, executive complexity, rhythmic complexity, number of notes played by the left hand, and number of notes played by the right hand. For each metric, mean values were calculated across corresponding excerpts by the two composers. Given the small sample size and the potential for non-normal distributions, we used two complementary non-parametric tests: the Sign Test, which evaluates the consistency in directionality of paired differences, and the Wilcoxon signed-rank test, which additionally considers the magnitude of the differences. All tests were two-tailed and statistical significance was set at α = 0.05.

The timing of individual quarter notes (2/2 time “cut time” for Bach; and 6/4 for Chopin) underwent repetead-measures ANOVAs with 2 factors of variability (“beat”, with 2 levels: *downbeat* and *upbeat*) and “subdivision” (2 for Bach and 3 for Chopin). Multiple post-hoc comparisons of means were performed by means of the Tukey tests. The Greenhouse-Geisser correction was also applied to compensate for possible violations of the sphericity assumption associated with factors having more than two levels. In this case, the modified degrees of freedom are reported together with the epsilon (ε) and the corrected probability level.

## Results

Number of gestures per measure and per second, melodic/harmonic, executive and rhythmic complexity was higher in Chopin than in Bach. Both a Sign test and a Wilcoxon signed-rank test indicated a significant difference in structural complexity between Chopin and Bach compositions (Sign test: Z = 2.27, p = 0.023; Wilcoxon: Z = 2.37, p = 0.018), consistently pointing to greater complexity in Chopin’s works. Hand asymmetry (in favour of the right hand) was similar in the 2 pieces (see Table 2). Beat tracking (tactus instability) revealed a much regular timing for Bach (89.3 BPM, SD= 9.18), than Chopin (154.74 BPM, SD= 48.8) because of differences in the pieces’ interpretation and style (Fig. 4). Both datasets deviated from a normal distribution and exhibited significantly different variances. Non-parametric Mann-Whitney and Levene tests revealed significant differences (p<0.00001).

**Table 2.** Complexity was assessed by two external professional pianists using a five-point Likert scale (1 = Low, 3 = Medium, 5 = High). The number of notes and gestures refers to the mean count per measure. Gestures were quantified independently of notes and sounds, taking into account the motor actions required, executed simultaneously by both hands. Mean gesture frequency (gestures per second) was calculated by dividing the total number of gestures per minute by 60. * Bars 65 and 125 in Chopin were incomplete and thus excluded.

| Bach | Parameters | Chopin |
| --- | --- | --- |
| 5.08 (SD = 2.38) | Mean Left-Hand Note Count per Bar | 8.37 (SD = 2.92) |
| 6.44 (SD = 2.3) | Mean Right-Hand Note Count per Bar | 9.46 (SD = 5.83) |
| 7.24 (SD = 1.74) | Mean Key-Onset Density per Bar | 8.15 (SD = 2.77) |
| 2.03 (SD = 0.84) | Mean Gesture Frequency (Hz) | 3.06 (SD = 1.07) |
| 3.27 (SD = 0.86) | Melodic/harmonic Complexity | 3.61 (SD = 1.43) |
| 3.36 (SD = 1.14) | Executive Complexity | 3.86 (SD = 1.14) |
| 2.88 (SD = 0.79) | Rhythmic Complexity | 3.44 (SD = 0.79) |
| 0.69 (SD = 0.14) | Average Quarter Note Duration (sec) | 0.46 (SD = 0.23) |
| 2.76 (SD = 0.51) | Average Bar Duration (sec) | 2.74 (SD = 1.09) |

**Fig. 4.**
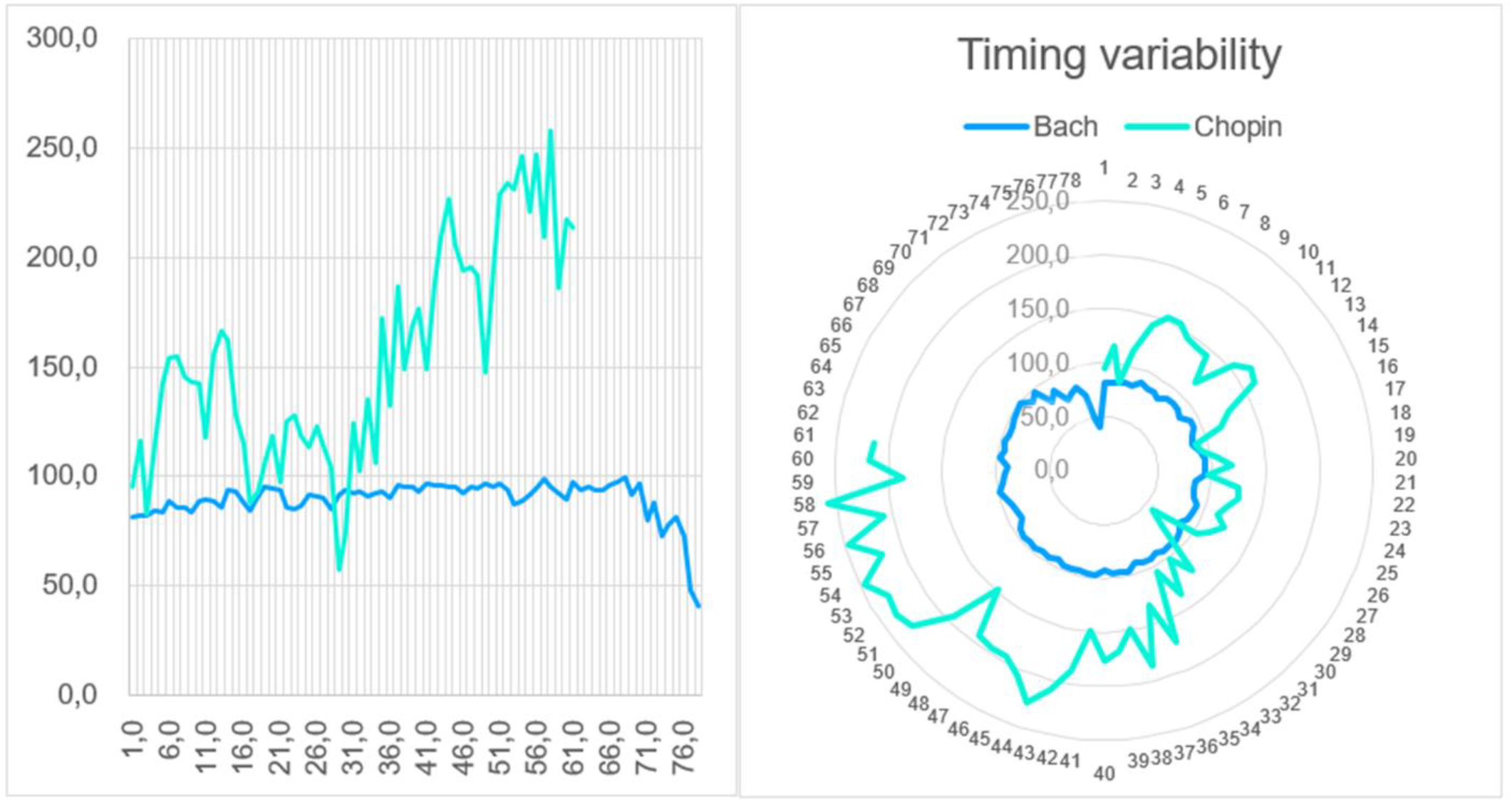
Temporal variability (expressed in BPM on the Y-axis) with respect to performances of Bach and Chopin. The X-axis represents the chronological order of bars within each execution (ranging from bar 1 to 78 in the Bach excerpt and from bar 65 to 125 in the Chopin excerpt).

Two-way repeated-measure ANOVAs revealed a significant interaction of “beat” x “subdivision” in tactus instability data for both Bach (F= 31.9, p < 0.00001) and Chopin (F= 33.7, p < 0.001; ε = 0.978, correct. p value = 0.0001). Tukey post-hoc comparisons (p<0.001) showed that quarters timing were significantly slower in the first subdivision in *downbeat* and in the last subdivision in *upbeat*, in both executions (see Fig. 5).

**Fig. 5.**
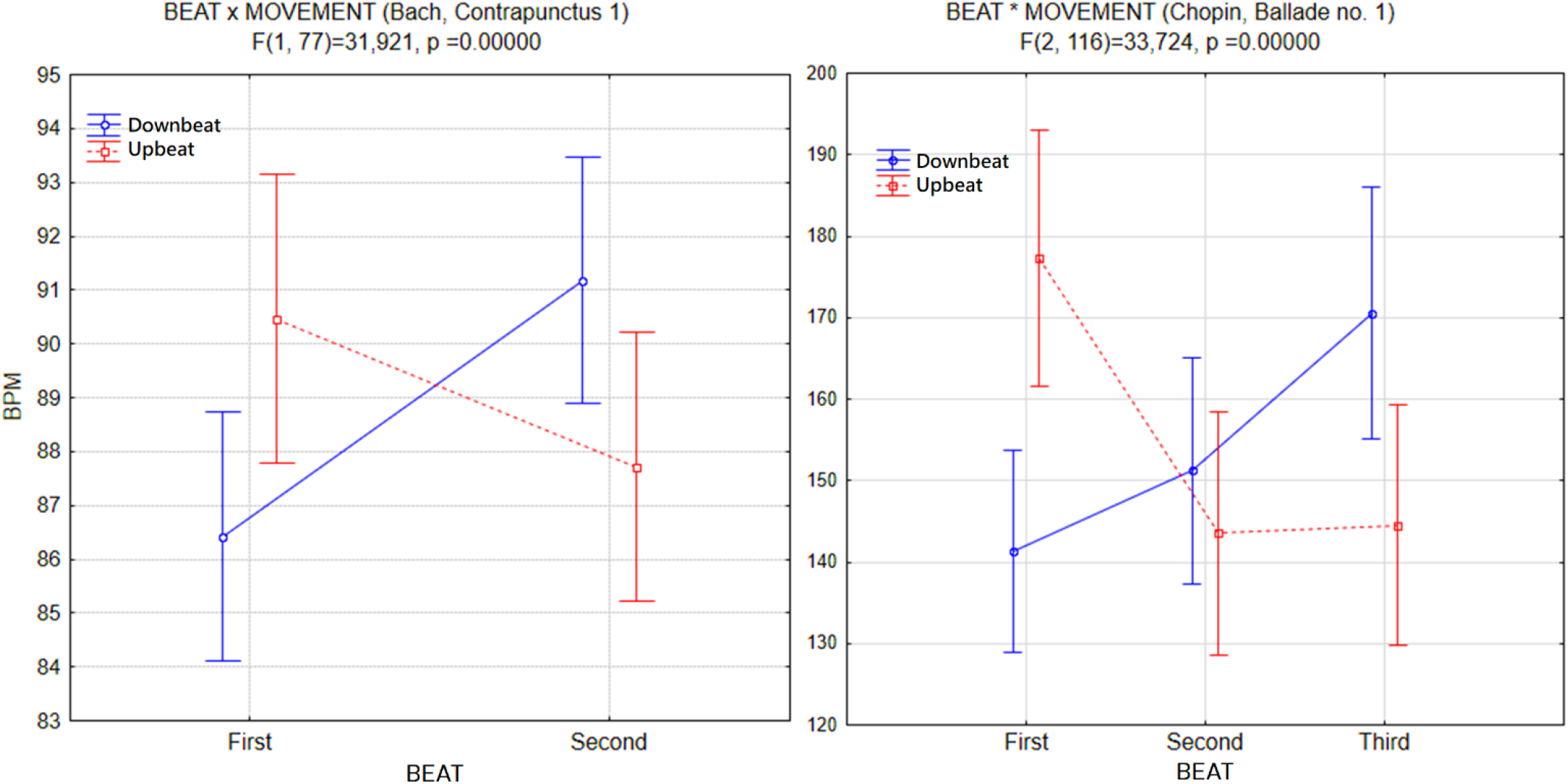
Interaction effects between beat type (Downbeat vs. Upbeat) and movement sections in two musical excerpts: Contrapunctus 1 by J.S. Bach (left panel) and Ballade no. 1 by F. Chopin (right panel). The plots display mean tempo values (in beats per minute, BPM) across different movement segments. Blue solid lines represent downbeat, while red dashed lines represent upbeat). Error bars indicate standard deviations. Significant BEAT × MOVEMENT interactions were observed for both excerpts indicating distinct temporal patterns across beats and musical subdivisions.

To derive the oscillation frequencies reported (≈ 0.362 Hz for Bach and ≈ 0.365 Hz for Chopin), we analyzed the periodic fluctuations in tactus inaccuracy, as expressed in beats per minute (BPM), across successive musical movements. Tactus inaccuarcy was operationalized as the variability in performed beat timing, reflecting micro-rhythmic deviations from an idealized metrical pulse. We applied time-series analysis to the BPM data, specifically focusing on recurring temporal patterns. The resulting wave-like structures revealed consistent oscillatory dynamics, with frequencies centered around 0.36 Hz for both excerpts (Fig. 6). These values indicate a stable rhythm in the fluctuation of timing precision, suggesting the presence of low-frequency periodic modulations in performers’ temporal control over extended musical spans.

**Fig. 6.**
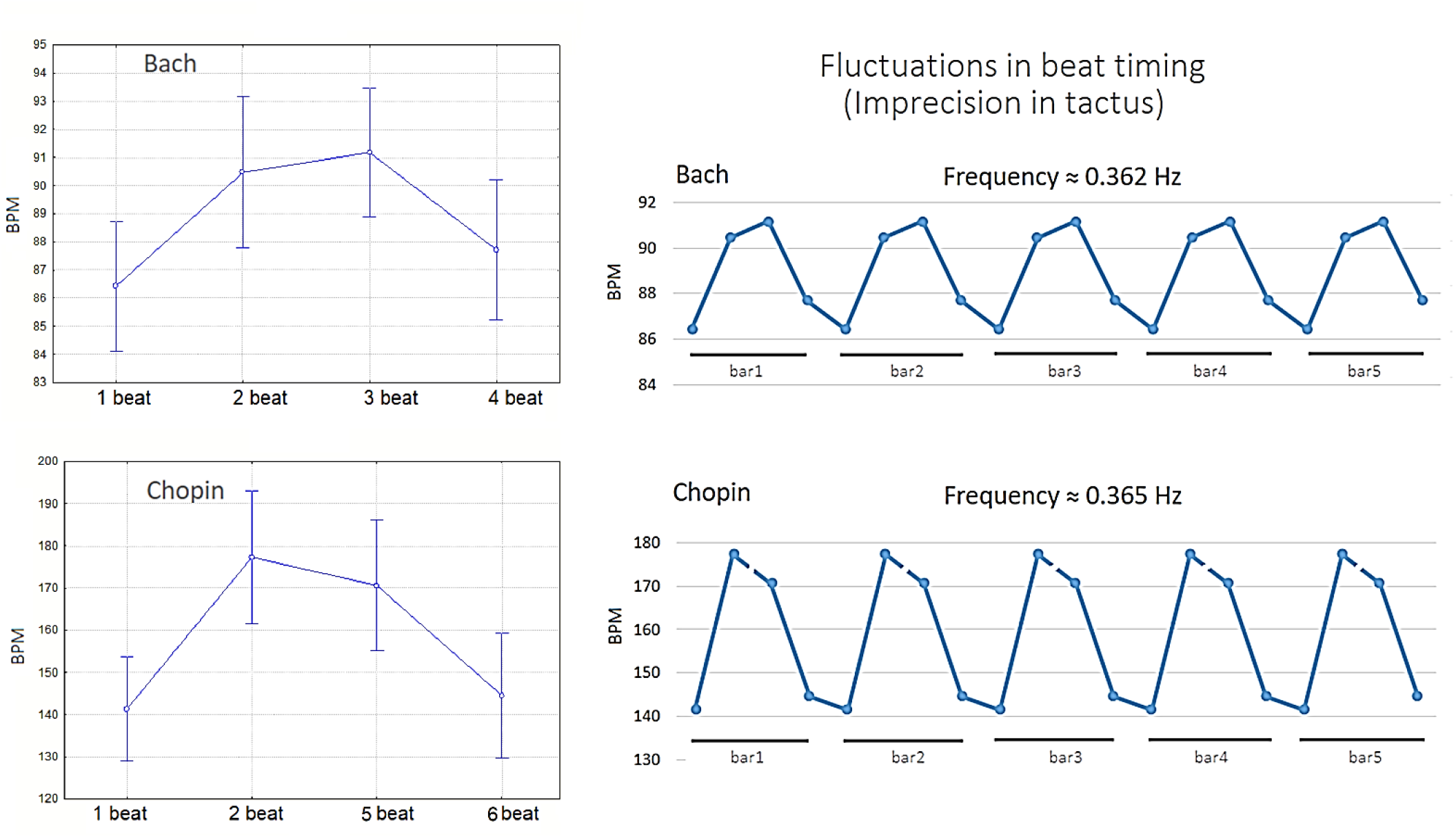
Oscillatory rhythm of tactus instability expressed in BPM for Contrapunctus I by J.S. Bach (top) and Ballade No. 1 by F. Chopin (bottom), focusing on the first and last beats. The analysis reveals low-frequency wave-like fluctuations in beat-level temporal variability, with an estimated oscillation frequency of approximately 0.362 Hz for Bach and 0.365 Hz for Chopin. These results suggest a periodic modulation of beat imprecision, indicative of structured temporal dynamics in performance.

## Discussion

As well known, Bach’s Baroque style feature regular emotional fluctuations and stable expressive changes, while Chopin’s Romantic composition highlights dramatic, free emotional expression with flexible rhythms (Anderson and Schutz, 2021). Our data revealed that the number of musical gestures per bar (as notated in the score) and per second (as executed in performance), as well as the estimated melodic/harmonic, executive, and rhythmic complexity—and particularly beat irregularity (i.e., imprecision in the tactus)—were all higher in Chopin compared to Bach. Interestingly, despite huge stylistic and interpretative differences, both performances revealed a shared underlying motor control structure with a similar periodic oscillatory pace of deviations from tactus. Time signature detection revealed a similar periodic oscillatory structure in the 2 pieces, with wavelength determined by the first downbeat’s duration. This increased progressively, peaked, and tapered with the final upbeat, forming a dynamic oscillatory pattern. Such periodicity may reflect intrinsic neural motor control and time signaling, as in speech and locomotion (e.g., central pattern generators). This framework might stabilize temporal coherence under varying expressive demands. The present analysis reveals that the tempo fluctuations in both Bach’s Contrapunctus 1 and Chopin’s Ballade No. *1* exhibit oscillatory patterns within the infra-delta frequency range (∼ 0.36 Hz). This frequency aligns with the lower spectrum of neural oscillations associated with meditative states, concentration, motor control and inhibition (e.g., Harmony et al., 2013) suggesting a potential entrainment between musical tempo and endogenous brain rhythms.

Neural entrainment, the synchronization of brainwave frequencies to external rhythmic stimuli, has been extensively documented in the context of auditory processing (Nozaradan, 2014; Aparicio-Terrés et al., 2025). For instance, studies have demonstrated that rhythmic auditory stimuli can modulate neural oscillations, leading to enhanced cognitive and perceptual outcomes. The function of music rhythmic entrainment in motor rehabilitative training and learning is long known (Thaut et al., 2015). Furthermore, the phenomenon of beat induction—the cognitive ability to perceive and synchronize with a beat—is considered fundamental to human music perception and is closely linked to neural entrainment mechanisms. These findings underscore the intricate relationship between musical structure and neural dynamics, highlighting the potential of music to engage and modulate brain rhythms.

The present findings lend support to the hypothesis that expressive piano performance—despite stylistic, structural, and historical variability—relies on a shared periodic motor scaffolding that may reflect intrinsic neural timing mechanisms. This account converges with, and sharpens, prior theoretical formulations (Friberg and Sundberg, 1999; Todd, 1992; Clarke, 1988), which posit an endogenously generated, hierarchically organized temporal reference signal closely aligned with the tactus that serves as a latent dynamical prior for action–perception coupling, constraining and parameterizing expressive microtiming through predictive sensorimotor processes and entrainment-based coordination (Madison, 2024). The observed regularities in tactus inaccuracy, with their oscillatory modulations peaking and tapering in synchrony with phrase structure, appear consistent with contemporary models of temporal encoding in distributed neural populations (Buonomano & Laje, 2010; Zhou & Buonomano, 2022).

Interestingly, recent evidence suggests that neural oscillations in delta and beta frequency bands serve as temporal integrators in sensorimotor synchronization, facilitating precise coordination between auditory perception and motor execution (Keitel et al., 2017; Arnal et al., 2015). In particular, delta-band entrainment has been implicated in predictive timing processes during musical performance, aligning internal rhythmic templates with external auditory cues (Nacher et al., 2013). Motor delta oscillations encode neural motor trajectories and are visible in the dynamics of most basic motor acts (Morillon et al., 2019). For example delta-band oscillations in motor regions predict hand selection for reaching Hamel-Thibault et al., (2018) and indeed phase-locked neural oscillations in the motor cortex are a prerequisite for the preparation and execution of motor actions (Popovych et al., 2016). Such entrainment is enhanced in expert musicians (Wang et al., 2022; Proverbio and Valtolina, 2025), who exhibit superior coherence in delta oscillations across the premotor cortex and parietal regions (Nandi et al., 2023). Other evidences indicate how oscillatory activity in the delta band aids rhythmic coordination during stationary locomotor behaviors (Fortunato et al., 2022). This neural architecture may underlie the temporal stability observed in the piano performances observed, even in the context of rubato and expressive timing deviations characteristic of Romantic repertoire.

Of particular interest is the apparent neurobehavioral parallel between rhythmic motor coordination in piano performance and other biologically rooted temporal behaviors, such as gait and speech. These behaviors, too, are governed by central pattern generators (CPGs), neural circuits capable of producing rhythmic output in the absence of sensory feedback (Mauk & Buonomano, 2004). In this vein, Klimesch (2018) advances the notion of a unified frequency architecture governing brain–body dynamics, whereby neural oscillations are organized in discrete, harmonically related bands that scale across temporal domains. Crucially, this framework posits a systematic cross-frequency coupling between slower peripheral rhythms (e.g., respiration and locomotor cycles) and faster cortical oscillations, resulting in a hierarchically nested structure. Such an organization may enable temporally precise coordination by constraining sensorimotor integration and facilitating predictive timing mechanisms, thereby providing a common dynamical substrate for both motor execution and higher-order cognitive functions. The emergence of a low-frequency oscillatory structure in both Bach and Chopin—despite their markedly differing rhythmic and even beat imprecision profiles—may thus reflect a conserved temporal architecture that scaffolds motor execution in humans. Finally, recent research in neuroplasticity underscores the transformative effect of musical training on the sensorimotor system, revealing enhanced connectivity within cortical-striatal-cerebellar loops (Herholz & Zatorre, 2012). These adaptations likely afford expert pianists the capacity to maintain internal rhythmic coherence even when expressive liberties are taken, as seen in the divergent but structurally comparable *tactus* patterns elicited by the two pieces analyzed here. Interestingly, a relationship between midfrontal delta-band (1–4 Hz) oscillations and interval timing in Parkinson’s disease patients experiencing freezing of gait was reported (Bosch et al., 2022). The findings suggest that impaired delta oscillations are associated with both cognitive dysfunction and gait abnormalities, highlighting the role of delta rhythms in coordinating motor and cognitive functions.

Together, these findings suggest that piano musical performance is governed by a complex interplay between hierarchical motor planning, predictive timing, and neural oscillations. The convergence of temporally structured deviations from *tactus* across pieces may serve as a behavioral signature of the neural dynamics underpinning high-level motor control in music. While the present study offers valuable insights from a single elite pianist, further research—including kinematic and EEG investigations across diverse performers and instruments—will be essential to substantiate and generalize these findings.

### Limitations

The present study is based on a single elite performer and should therefore be considered a proof-of-concept investigation rather than a generalizable account of musical timing. While the observed oscillatory patterns are consistent with theoretical models of neural timing, no direct electrophysiological evidence is provided to establish a causal link between behavioral dynamics and neural oscillations. Future research combining kinematic, behavioral, and electrophysiological measures across multiple performers will be necessary to validate and extend these findings.

## Acknowledgments

We extend our heartfelt gratitude to the virtuoso pianist ‘P1’ for his irreplaceable collaboration and artistic insight.

## Ethical approval and informed consent statements

The participant provided written informed consent prior to participation. The study procedures were conducted in full accordance with the principles outlined in the Declaration of Helsinki and received formal approval from the competent institutional ethics committee (RM-2023-605, 01/10/2023).

## Data availability statement

The datasets generated during the current study is stored in this repository: Qin, Chang; Proverbio, Alice Mado (2025), “Data for “Infra-Delta Oscillatory Signatures and Gesture Density in Expert Piano Performance”“, Bicocca Open Archive Research Data, V1, doi: 10.17632/yp3kgbtz7g.1

